# The nuclear receptor NR4A1 is regulated by SUMO modification to induce autophagic cell death

**DOI:** 10.1101/745026

**Authors:** Gabriela Zárraga-Granados, Gabriel Muciño-Hernández, María R. Sánchez-Carbente, Wendy Villamizar-Gálvez, Ana Peñas-Rincón, Cristian Arredondo, María E. Andrés, Christopher Wood, Luis Covarrubias, Susana Castro-Obregón

**Affiliations:** Departamento de Neurodesarrollo y Fisiología, División de Neurociencias, Instituto de Fisiología Celular, Universidad Nacional Autónoma de México; Biotechnology Research Center, Universidad Autónoma del Estado de Morelos. Av. Universidad 1001, Chamilpa, Cuernavaca, Morelos, México 62209; Departamento de Biología Celular y Molecular, Facultad de Ciencias Biológicas, Pontificia Universidad Católica de Chile; Laboratorio Nacional de Microscopía Avanzada, Instituto de Biotecnología, UNAM; Departamento de Genética del Desarrollo y Fisiología Molecular, Instituto de Biotecnología, UNAM

**Author notes:** **Corresponding author:** (SCO).

**Keywords:** NR4A1, SUMO, autophagy, cell death, Substance P, NK_1_R

## Abstract

NR4A is a nuclear receptor protein family whose members act as sensors of cellular environment and regulate multiple processes such as metabolism, proliferation, migration, apoptosis, and autophagy. Since the ligand binding domains of these receptors have no cavity for ligand interaction, their function is most likely regulated by protein abundance and post-translational modifications. In particular, NR4A1 is regulated by protein abundance, phosphorylation, and subcellular distribution (nuclear-cytoplasmic translocation), and acts both as a transcription factor and as a regulator of other interacting proteins. SUMOylation is a post-translational modification that can affect protein stability, transcriptional activity, alter protein-protein interactions and modify intracellular localization of target proteins. In the present study we evaluated the role of SUMOylation as a posttranslational modification that can regulate the activity of NR4A1 to induce autophagy-dependent cell death. We focused on a model potentially relevant for neuronal cell death and demonstrated that NR4A1 needs to be SUMOylated to induce autophagic cell death. We observed that a triple mutant in SUMOylation sites has reduced SUMOylation, increased transcriptional activity, altered intracellular distribution, and more importantly, its ability to induce autophagic cell death is impaired.

**Summary Statement:** The modification of the nuclear receptor NR4A1 by SUMO regulates its transcriptional activity, intracellular localization and is required to induce autophagic cell death.

## Introduction

Nuclear receptors are a superfamily of transcription factors involved in a vast number of biological processes. They share three common structural domains: a N-terminal transactivation domain (TAD), a central double zinc finger DNA binding domain (DBD) and a C-terminal ligand binding domain (LBD). Among the known human nuclear receptors, the ones belonging to the NR4A family act as sensors of the cellular environment and contribute to cell fate decisions, such as cell proliferation, differentiation, migration, cell death, etc. Physiologically, NR4A members influence the adaptive and innate immune system, angiogenesis, metabolism and brain function. The NR4A family is comprised of three members that bind the same DNA elements (1): NR4A1 (Nur77, TR3, NGF1B, etc.), NR4A2 (Nurr1, NOT, TINUR, etc.) and NR4A3 (Nor1, MINOR, etc.). NR4A family members are considered orphan nuclear receptors since endogenous ligands are unknown and, although several natural and artificial compounds enhance their transcriptional activity, they are able to activate transcription by solely up-regulating the expression of their genes (1). Therefore, rather than regulation by ligand interactions, their endogenous function is most likely regulated by NR4A family members protein abundance and post-translational modifications (PTM). All NR4A family members are mainly modified by phosphorylation by over 20 different kinases described so far, and at least NR4A1 is also acetylated by CBP/p300 and deacetylated by HDAC1. Specific PTM affect their interaction with either DNA or other proteins, as well as their intracellular localization (1).

A wide variety of stimuli induces the expression of *Nr4a* genes, both during development and in adulthood. For example, they are induced in response to caloric restriction (2), exercise (3) or during learning and long term memory (4, 5). NR4A family proteins regulate the expression of several genes, some of which are involved in lipid and glucose metabolism, insulin sensitivity and energy balance. Another important function of NR4A family proteins is to prevent DNA damage and to promote DNA repair (6, 7). NR4A1 also has non-genomic activities both in the nucleus and in the cytoplasm, altering the function of interacting proteins. For all the functions described above, understanding the molecular regulation of NR4A activity is an active area of research.

Autophagy is mainly a catabolic process that allows cells to recycle macromolecules when needed or to eliminate damaged proteins and organelles, among other components, contributing to cell health (8); occasionally, however, it also contributes to an alternative secretion mechanism (9) or even to cell death (10). When inhibition of autophagy prevents cell death, it is referred to as autophagic cell death, although the actual cause of cell death still needs to be understood. We found previously that NR4A1 plays an essential role in a form of cell death induced by several stimuli that is non-apoptotic (in apoptosis-competent cells) and is dependent on autophagy. NR4A1 inactivation by either over-expression of dominant negative mutants or by RNAi prevents cell death, and inhibition of autophagy either pharmacologically or by RNAi to reduce autophagic gene expression also prevents cell death (11–13). Hence, NR4A1 mediates autophagic cell death, a phenomenon that was subsequently confirmed by others. In the case of melanoma cells treated with THPN, a chemical compound targeting NR4A1, cell death occurs after induction of excessive mitophagy, due to NR4A1 translocation to the inner membrane of the mitochondria, causing dissipation of mitochondrial membrane potential by the permeability pore complex ANT1/VDAC (14). Furthermore, Dendrogenin A, a mammalian cholesterol metabolite ligand of liver-X-receptors (LCRs) induces *Nr4a1* expression in association with excessive autophagic cell death both *in vitro* and *in vivo* (15, 16), displaying anticancer and chemopreventive properties in mice(17). Interestingly, NR4A1 interaction with anti-apoptotic BCL2 family members outside the BH3 domain induces autophagic cell death (18), which seems to be mediated by releasing BECN1 in a model of cigarette smoke-induced autophagic cell death(19). Taken together, these findings indicate that NR4A1 could function as a broad inducer of autophagic cell death. Some authors have coined autophagic cell death as autosis (20); we will use this term hereafter for simplicity, although whether NR4A1-induced autophagic cell death is mediated by Na^+^,K^+^-ATPase pump, as has been documented for autosis, has not been addressed.

In the present work we aimed to study whether specific PTM confer upon NR4A1 the ability to induce autosis, by using our already described model of NR4A1-mediated autosis, induced upon Substance P (SP) binding to its NK1R receptor (here referred to as SP/NK1R-induced autosis). SP is a neuropeptide involved in several physiological functions and pathological situations, including emotional behavior, pain perception, addiction, inflammation, neurodegeneration, etc. Accordingly, NK1R is expressed all around the body, including endothelial cells and in both central and peripheral nervous system, and SP is present in all body fluids such as cerebrospinal fluid, blood, etc. (21). We focused in this model for its relevance in neuronal cell death, as interfering with SP signaling reduces infarct volume and neuronal cell death in *in vivo* models of ischaemia (22) and of excitotoxin-induced seizures (23), both situations in which NR4A1 expression is also induced *(i.e*. ischaemia (24) and kainic acid-triggered seizures (25)).

During SP/NK_1_R-induced autosis, NR4A1 is regulated by both protein abundance and phosphorylation (12), and undergoes nuclear-cytoplasmic translocation (13), potentially having both genomic and non-genomic functions. A previous report showed that NR4A1 undergoes sequential SUMOylation and ubiquitination, which together control the degradation of NR4A1 after induced stress (26). SUMO modification can affect protein stability, transcriptional activity, protein-protein interactions and intracellular localization of target proteins (27). In addition, SUMO modification affects numerous cellular processes overlapping with those described for NR4A1 function, both in development and adulthood. Even though the SUMO machinery is mainly located in the nucleus, numerous cytoplasmic proteins have also been identified to have their function modulated by SUMO modification (28, 29). Therefore, we hypothesize that NR4A1 might be SUMOylated in response to SP/NK_1_R signaling, conferring upon it the ability to induce autosis.

SUMOs are small ubiquitin-related peptides, approximately 11 KDa size, that become conjugated to a variety of proteins and are deconjugated by SUMO-specific proteases. Despite the name, SUMO shares less than 20% homology with ubiquitin. The similarity is more significant in the biochemical mechanism of ligation, as it involves three conjugating enzymes: E1, a dimer known as SAE1/SAE2; E2, a protein named UBC9; and E3, which are a group of proteins conferring target specificity. There are four genes in mammals coding for four SUMO peptides (SUMO1-4). SUMO2 and 3 share 95% homology and there are no antibodies that distinguish between them, so they are referred to as SUMO2/3; they are only 50% identical to SUMO1 (reviewed in (30)).

NR4A1 is an early response gene, whose expression is induced in minutes, and the protein is degraded after a couple of hours. However, we observed that NR4A1 expression is sustained during SP/NK_1_R–induced autosis, suggesting that in this scenario NR4A1 protein would scape degradation. In this work, we confirmed that NR4A1 is indeed SUMOylated during SP/NK_1_R–induced autosis, and mutants with reduced SUMOylation showed increased transcriptional activity and altered intracellular distribution. More importantly, the ability to promote SP/NK_1_R–induced autosis was impaired when SUMOylation of NR4A1 was reduced.

## Experimental Methods

### Cell culture, plasmid transfection and cell death evaluation

Human embryonic kidney 293 (HEK293) cells were grown in high glucose DMEM (Invitrogen, Carlsbad, CA) supplemented with 10% fetal bovine serum (Sigma, St. Louis, MO) and penicillin/streptomycin 100 U/ml (Invitrogen, Carlsbad, CA). The cultures were incubated at 37°C in 95% air and 5% carbon dioxide with 95% humidity. Transient transfection was performed with Polyethylenimine (PEI, 25 kDa Polysciences Inc. # 23966-2) mixed with DNA in a 3:1 ratio. Briefly, 2×10^5^ cells/well were seeded into 35 mm wells 16-20 hr prior to transfection. Transfection solution: 1 μg of DNA was diluted into 75 μl OPTIMEM; 3 μg PEI was diluted into 75 μl OPTIMEM; then mixed and incubated for 15 minutes at room temperature. 0.5 ml of medium was removed from each well and the mixture was added dropwise to the cells. After 4 hr at 37 °C the medium was refresh supplemented without antibiotics. After 24 hr, 100 nM SP (SIGMA) was added when necessary. Expression of each construct in the transient transfections was determined by Western blot or immunofluorescence. Transient transfection efficiencies were in all cases >80%. The plasmid pcDNA3.1-N_1_R has previously been described (11, 31). Dr. Jacques Drouin (Laboratoire de Génétique Moléculaire, Institut de Recherches Cliniques de Montréal, Canada) kindly provided POMC-Luc, NBRE-POMC-Luc and NuRE-POMC-Luc reporter plasmids. Site-directed mutagenesis was performed using QuickChange Kit (Invitrogene). The alignment and prediction of consensus motifs was performed using Clustal W2 software, using the sequence UniProtKB - P22736 (NR4A1_HUMAN). All position information refers to this entry. Cell death was determined by Trypan blue exclusion or LDH release. The software PRISM 6.0 (GraphPad Software, La Jolla, CA, USA) was used for the one-way ANOVA statistical analysis, and the p values between indicated treatments in figures were calculated by Bonferroni’s Multiple Comparison Test.

### Western blot, immunoprecipitation and immunofluorescence analysis

For Western blotting, the transfected human embryonic kidney 293 cells were washed with cold PBS and homogenized in lysis buffer (150 μM NaCl, 1% Triton X-100, 50 μM Tris HCl pH 8.0, proteinases inhibitor cocktail cOmplete ULTRA and phosphatases inhibitor cocktail PhosSTOP (Roche Diagnostic Corporation, Indianapolis, IN, USA)). Cytoplasmic extracts were collected after 10 min. centrifugation at 14,000 rcf. Protein was quantified by Bradford assay and electrophoresis of equal amounts of total protein was performed on SDS-polyacrylamide gels. Separated proteins were transferred to polyvinylidene fluoride membranes at 4° C. Membranes were probed with the following antibodies: SUMO1 (1:1000 #4930), SUMO2/3 (1:1000 #4971), phospho-Threonine (1:500 #9381), Myc (1:350 #2276) and NR4A1 (1:2000 #3960) from Cell Signaling Technology Inc., Danvers, MA, USA; GAPDH (1:8000, Research Diagnostics, Flanders, NJ, USA); TUBULIN (1:7000, Abcam, Cambridge, MA, USA); FLAG (1:1000, #F3165 SIGMA, San Luis, Missouri, USA); UBC9 (1:250, #610748 BD Biosciences, San Jose, CA); UBIQUITIN (1:1000, #U5379 SIGMA, San Luis, Missouri, USA); HA (#H6908 SIGMA, San Luis, Missouri, USA). The membranes were incubated in the appropriate horseradish peroxidase-coupled secondary antibody for 1 hr followed by enhanced chemiluminescence detection (Amersham, Arlington Heights, IL). Alternatively, appropriate infrared dye-coupled secondary antibodies (1:10,000 dilution of anti-rabbit IRDye800 and anti-mouse IRDye700, Rockland, Gilbertsville, PA, USA) were used and the blots were scanned in an Odyssey Imager (LI-COR Biosciences, Lincoln, Nebraska, USA). The immunoprecipitations were carried out using super-paramagnetic Microbeads conjugated to protein A or protein G, following the manufacturer’s instructions (MACS; Milteyi Biotec, Auburn, CA). For immunofluorescence, cells were seeded into Lab-Tek CC2 treated slide chambers (Nalgene Nunc International, Napeville, IL, USA); after washing with PBS cells were fixed with 4% paraformaldehyde/PBS for 10 min; then washed with PBS and permeabilized with 0.2% Triton X-100/PBS for 10 min. Afterwards cells were washed with PBS, pre-incubated 30 min with 4% BSA/PBS and 4% goat serum and washed again with PBS. Anti-NR4A1 (M-210 Santa Cruz Biotechnology, Santa Cruz, CA, USA) was diluted 1:200 in 2%BSA/PBS and incubated over night at 4°C. After washing with PBS cells were incubated with antirabbit coupled to Alexafluor 594 (Invitrogen, Carlsbad, CA) diluted 1:1000 in 2%BSA/PBS for 30 min at room temperature. Then, cells were treated with 10mg/ml RNAse for 30 min at 37°C and washed again with PBS. Cells were counterstained with 10 ng/ml DAPI and mounted for microscope observation on an Axiovert 200M (Carl Zeiss) confocal microscope.

### RNAi

Four regions were targeted for 3’UTR hNR4A1 mRNA (GenBank NM_002135) starting at positions: 2383 5’gcgccgugcuguaaauaaguu3’; 2401 5’ gcccagugcugcuguaaauuu3’; 2529 5’ccacauguacauaaacuguuu3’; 2535 5’guacauaaacugucacucuuu3’. The corresponding siRNAs were simultaneously transfected. These siRNAs were purchased as SMARTpool from Dharmacon (Lafayette, CO, USA). Control siRNA targeting a protein of Rotavirus was a kind gift from Dr. Susana López (Instituto de Biotecnología, UNAM).

siRNA transfection: Human embryonic kidney 293 cells (10^5^ cells per well in 12-well plates) were grown in high-glucose DMEM supplemented with 10% fetal bovine serum (Sigma, St. Louis, MO), with no antibiotics for 16 hr. The siRNA specific for each target gene was transfected with Lipofectamine 2000 reagent (Invitrogen, Carsband, CA, USA) according to the manufacturer’s instructions, using 3 μg siRNA: 6 μl Lipofectamine 2000 ratio. After 4 hr of incubation, plasmids were transfected.

### Transcription Assays

Human embryonic kidney 293 cells were seeded in 24-well plates (1×10^5^ cells/well). Transient transfections were performed with 2.1 μg PEI plus 700 ng DNA [225 ng NK_1_R; 225 ng reporter plasmid (POMC minimal promoter-Luciferase that lacks responsive elements, NBRE-Luciferase or NurRe-Luciferase); 225 ng NR4A1 (or mutants) and 25 ng of a plasmid encoding Renilla luciferase to normalize transfections]. Twenty-four hours after transfection, the cells were incubated or not with 100 nM SP for 3 h. Luciferase and Renilla activities were determined using the Dual-Luciferase Reporter Assay (Promega #E1980), according to manufacturer’s instructions and a FLUOstar OMEGA luminometer (BMG LABTECH).

## Results

### NR4A1 is SUMOylated during SP/NK_1_R–induced autosis upon previous phosphorylation

SUMO peptides are conjugated to a Lys residue, frequently within a consensus motif ΨKXE, where Ψ represents any hydrophobic residue and X any amino acid(30). It has been shown that NR4A2 is SUMOylated in two motifs conserved among the family members that lead to SUMO ligation to lysines K91 (32) or K558 (33), depending on the cell context. In order to find additional potential SUMOylation sites also conserved in the three members of the family, we aligned NR4A protein sequences and searched for the SUMO consensus motif ΨKXE. We identified K102 and K577 within a SUMO motifs, which are indeed SUMOylated (26), and observed that K558 is conserved in NR4A1 and NR4A2, and hence it is also a potential site for SUMOylation in NR4A1 (positions numbers refer to the human canonic sequence UniProtKB P22736) (Figure 1A).

**Figure 1.**
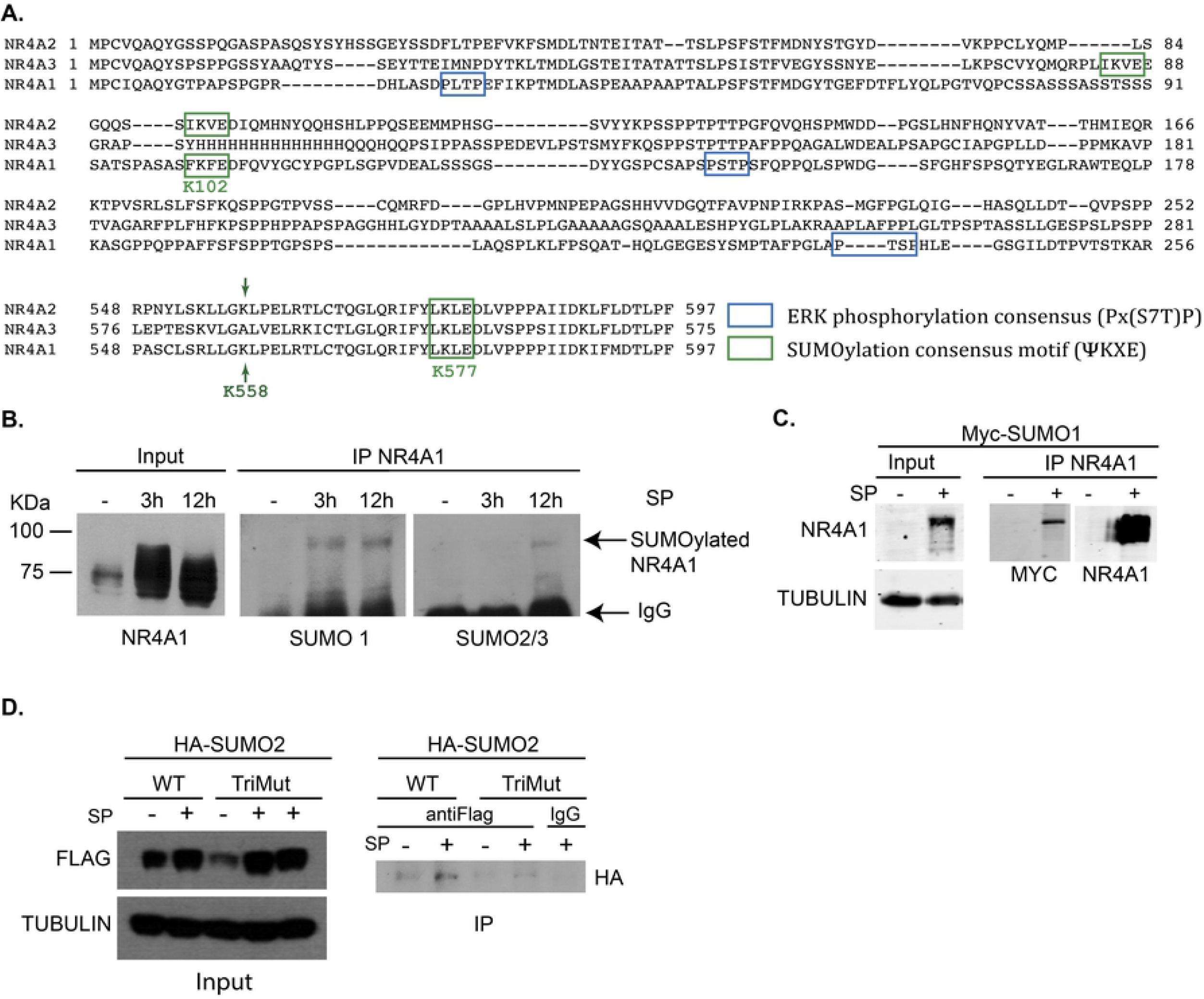
NR4A1 is SUMOylated. **A**, *Protein sequence alignment of NR4A family members*. Two conserved SUMOylation consensus motifs among members (green squares) and three potential ERK phosphorylation consensus sites in NR4A1 (blue squares) are shown. K558 is conserved in NR4A2 and NR4A1. **B**, *NR4A1 is SUMOylated in response to SP*. Immunoprecipitation (IP) of NR4A1 from cells transfected with NK_1_R expression vector and exposed to SP for the indicated times, to detect by Western blot either SUMO1, SUMO2/3 or NR4A1. IgG from IP was detected by the secondary antibody. **C**, *Over-expression of myc-SUMO1 enhances the amount of NR4A1 conjugation to SUMO1 in response to SP*. Cells were transfected with NK_1_R and MYC-tagged SUMO1 expression vectors (Myc-SUMO1) and exposed (+) or not (-) to SP for 3 hr. NR4A1 was immunoprecipitated and developed by Western blot to detect Myc tag or NR4A1. Tubulin was detected in whole extracts as a loading reference (Input). **D**, *Triple mutant NR4A1_K102,558,577R (TriMut) has reduced SUMOylation*. Cells were transfected with expression vectors for HA tagged SUMO2, FLAG-NR4A1 wild type or FLAG-NR4A1_TriMut, and NK_1_R to allow post-translational modifications in response to SP (+). Immunoprecipitation was performed with FLAG antibody (antiFLAG) or IgG as a control of irrelevant antibody. WB was developed with antibodies against indicated proteins. TUBULIN was detected as a loading reference (Input).

To analyze NR4A1 SUMOylation during SP/NK_1_R–induced autosis, we immunoprecipitated NR4A1 at different time points after SP exposure and looked for SUMO1 or SUMO2/3 conjugation by Western blot. Starting at 3 hr after SP induction, a fraction of NR4A1 was SUMOylated by SUMO1 and the modification lasted up to 12 hr, a time point at which SUMO2/3 ligation was also observed (Figure 1B). We verified that SUMO peptide conjugation of NR4A1 occurs during SP/NK_1_R–induced autosis by overexpressing Myc-tagged SUMO1 (Figure 1C) or HA-tagged SUMO2 (Figure 1D). To study the role of SUMOylated NR4A1 we substituted the three lysine residues expected to be modified by SUMO (K102, K558 and K577) for arginine. First, we evaluated whether the triple NR4A1_K102,558,577R mutant (named TriMut) had reduced SUMOylation in comparison to the level determined in NR4A1 wild type. We immunoprecipitated FLAG-NR4A1 wild type or FLAG-NR4A1_TriMut and looked for the presence of SUMO2 by Western blot. Indeed, NR4A1_TriMut showed reduced SUMOylation (Figure 1D), although additional Lys residues seem to be also modified by SUMO, as a weak signal was still detected.

There are several examples of crosstalk between phosphorylation and SUMOylation, having phosphorylation both positive and negative regulation over SUMO modification (34). During SP/NK_1_R–induced autosis NR4A1 is phosphorylated by ERK2, which is necessary for autosis induction, as the inhibition of MEK2 and ERK2 (but not the inhibition of MEK1 or ERK1) prevents NR4A1 phosphorylation and SP/NK_1_R–induced autosis (12). Accordingly, NR4A1 is phosphorylated *in vitro* by ERK2 but not by ERK1 in threonine 143 (35). Therefore, we mutated T143 to alanine, expecting to abolish its phosphorylation. Indeed, T143 is phosphorylated during SP/NK_1_R–induced autosis, since mutant NR4A1_T143A had reduced phosphorylation level as compared to NR4A1 wild type (Figure 2A). Nevertheless, a weak signal was still observed, indicative of an additional phosphorylated threonine. We found two additional ERK consensus motifs in the NR4A1 sequence (Figure 1A), although additional phosphorylation sites could be targeted by other kinases.

**Figure 2.**
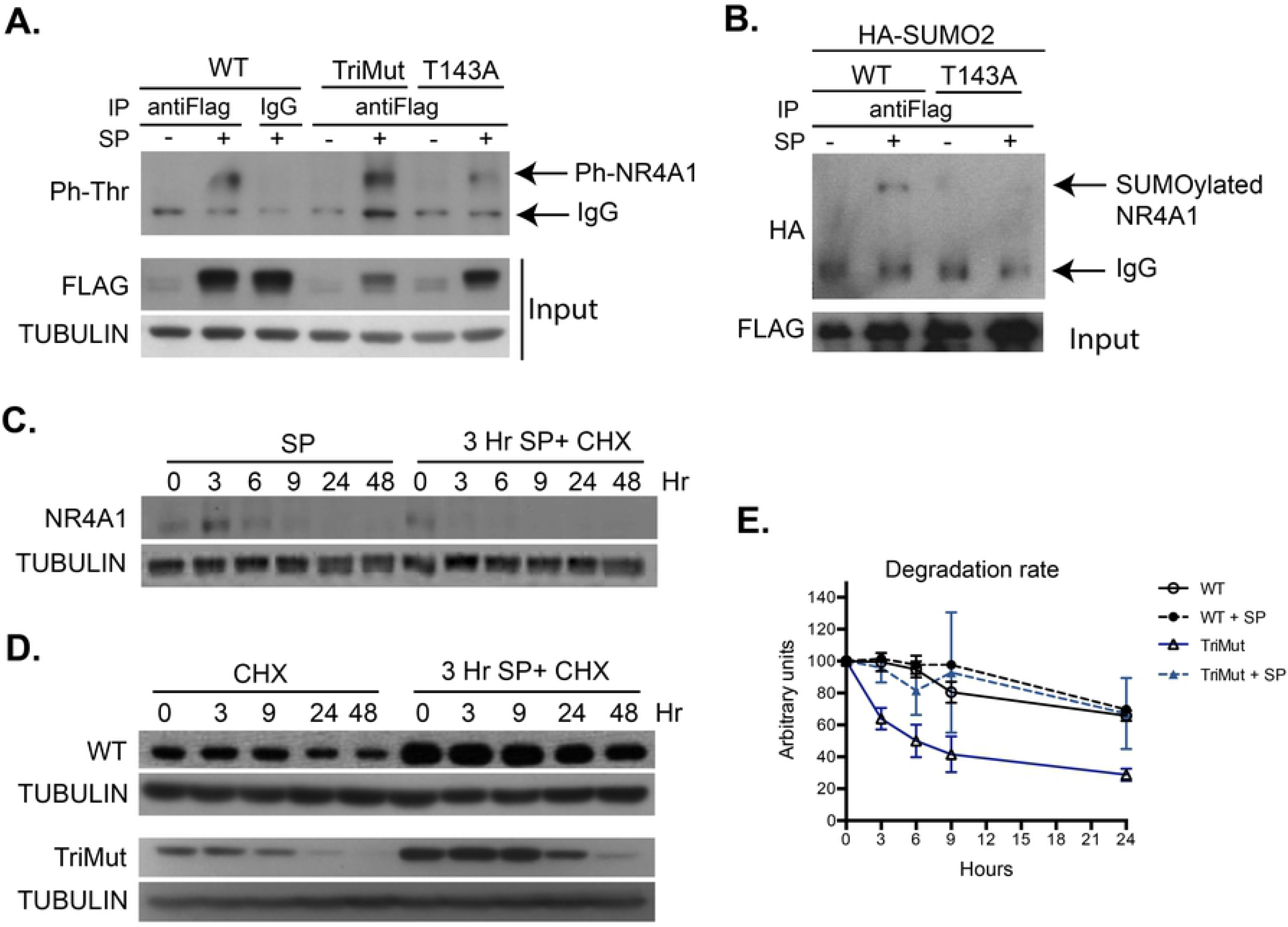
NR4A1 SUMOylation depends on former phosphorylation. **A**, *SUMOylation in K102, K558 and K577 is not necessary for NR4A1 phosphorylation in response to SP*. Cells were transfected with expression vectors for FLAG-NR4A1 wild type or FLAG-NR4A1_TriMut and NK_1_R, and exposed (+) or not (-) to SP for 3 hr. Total protein extracts were immunoprecipitated with anti-FLAG or IgG and the level of threonine phosphorylation was estimated by WB. TUBULIN was detected for comparison of total protein extraction for IP, and FLAG to show the level of expression of each construct. **B**, *Phosphorylation in T143 in necessary for NR4A1 SUMOylation in response to SP*. Cells were transfected with expression vectors for FLAG-NR4A1 wild type or FLAG-NR4A1_T143A, HA-SUMO2 and NK_1_R, and exposed (+) or not (-) to SP for 3 hr. Total protein extracts were immunoprecipitated with anti-FLAG and the level of SUMOylation was estimated by WB detecting HA. FLAG was detected to show the level of expression of NR4A1 constructs. **C**, *NR4A1 peak of synthesis is at 3 hr after SP addition and has a half-life of less than three hours*. Lanes 1-6, total protein extracts were obtained from cells transfected with NK_1_R and exposed to SP for the indicated times. Lanes 7-12, total protein extracts were obtained from cells transfected with NK1R expression vector, treated for 3 hr with SP to reach maximum expression of NR4A1 (considered time 0 for cycloheximide treatment), and then exposed to cycloheximide (CHX) for the indicated time to inhibit new protein synthesis. Three hr after CHX less than half of the initial amount of NR4A1 protein remains. **D**, *NR4A1 basal stability is enhanced by SUMOylation, but TriMut still becomes stabilized in response to SP signaling*. To compare stability of NR4A1 wild type with NR4A1_TriMut, cells were transfected with expression vectors for NK_1_R and either FLAG-NR4A1 or FLAG-NR4A1_TriMut. 24 hr after of transfection cells were treated with CHX for the indicated times, or treated for 3 hr with SP and then with CHX for the indicated time. Clearly, the amount of FLAG-NR4A1_TriMut was reduced and was degraded faster than wild type, but yet responded to SP induction. E. Quantitative comparison of the degradation rate of NR4A1 WT and TriMut, with or without SP, from densitometric analysis of at least three independent experiments described in D. Each blot was normalized with their corresponding TUBULIN. The measurement obtained for each NR4A1 (WT or TriMut) before CHX treatment (time zero) was arbitrary considered 100 units. Notice that the faster degradation rate of TriMut is overcome in response to SP. Each dot represents the mean and bars represent standard deviation.

To analyze whether SUMO conjugation influences phosphorylation, we studied the level of phosphorylation when SUMOylation is reduced. As shown in Figure 2A, the level of phosphorylation on threonine residues was not affected in NR4A1_TriMut. This result indicates that phosphorylation is not affected by SUMOylation in this case. On the other hand, SUMOylation resulted dependent on previous phosphorylation in T143, as NR4A1_T143A was barely SUMOylated (Figure 2B). Therefore, NR4A1 SUMOylation, subsequent to phosphorylation, could alter NR4A1 stability and/or intracellular distribution and, consequently, the ability to induce autosis.

In previous work we observed that *Nr4a1* expression is induced by SP signaling (12), but NR4A1 protein abundance cannot be explained by transcriptional regulation alone. We noticed that NR4A1 produced by expression of its cDNA (lacking both 5’ and 3’ UTR) from a viral promoter (CMV) that lacks endogenous regulatory regions, also accumulates in response to SP/NK_1_R (for example, look at Figure 2A, Input). We rationalized then that NR4A1 stability should be post-translationally regulated. MAPK signaling activated by SP/NK_1_R does not affect NR4A1 abundance, as it still accumulates when ERK2 signaling is inhibited (12). SUMO modification commonly enhances protein stability by binding to lysine residues that otherwise would be ubiquitinated, targeting the protein for proteasome degradation; nevertheless, it has also been observed that some E3 ubiquitin ligases, such as RNF4, which have both SIM (SUMO interacting motif) and RING domains, attach ubiquitin to SUMO-modified proteins (30), and this mechanism has been described to occur to NR4A1(26). Therefore, we asked whether NR4A1 SUMOylation regulated NR4A1 stability during SP/NK_1_R–induced autosis. First, we determined the endogenous NR4A1 half-life. NR4A1 reached a maximum level of expression 3 hr after SP exposure and by 9 hr it was still detected but clearly reduced (Figure 2C). We then inhibited new protein synthesis (with cycloheximide) after 3 hr of induction with SP (to start with the highest amount of NR4A1), and observed that NR4A1 was degraded very rapidly, as 3 hr after cycloheximide addition it was no longer detected (Figure 2C).

To compare the half-life of wild type NR4A1 with NR4A1_TriMut with reduced SUMOylation, we tagged it with FLAG, and compared FLAG-NR4A1 wild-type half-life with FLAG-NR4A1_TriMut (which allowed to distinguish NR4A1_TriMut from endogenously induced NR4A1). Since these constructs are expressed from a strong constitutive promoter (CMV), FLAG-NR4A1 started in these experiments with a higher amount of protein than the endogenously-induced NR4A1, therefore its expression could be observed it for a longer period of time. We inhibited new protein synthesis 24 hr after transfection. Notably, the stability imposed by SP signaling still occurred in FLAG-tagged NR4A1 (Figure 2D, upper panel). Under basal expression SUMOylation seemed to indeed contribute to NR4A1 stability, as the half-life of TriMut (which was SUMOylated to a lesser degree) was clearly reduced compared to wild type. Surprisingly, however, TriMut abundance still increased in response to SP (Figure 2D, bottom panel) reducing its degradation rate. Other reports describe that NR4A1 stability can be modulated by acetylation, which also occurs at Lys residues. We looked for acetylation during SP/NK_1_R–induced autosis, but no difference was found in the level of acetyl-Lysine content in immunoprecipitated NR4A1 from non-treated cells compared with SP treated cells (data not shown). We do not know at this point which mechanism mediates the increase in NR4A1 stability in response to SP signaling but, possibly, the remaining SUMOylation sites observed in TriMut (Figure 1D) could be involved.

### SUMOylation and phosphorylation alter NR4A1 transcriptional activity and intracellular distribution

SUMO modification of transcription factors usually inhibits their transcriptional activity, either by recruiting corepressors to promoter regions or by sequestering the SUMOylated form in nuclear bodies (36). NR4A1 transcriptional activity could also be negatively regulated by SUMOylation, since NR4A1 interacts with HDAC1, potentially recruiting corepressor complexes (1), and with PML, being potentially recruited into nuclear bodies (37). In previous studies we found that NR4A1 is transcriptionally active during SP/NK_1_R–induced autosis (13). Therefore we analyzed whether the transcriptional activity of NR4A1 is increased in the triple mutant in which predicted Lys targets of SUMOylation were replaced by arginine. NR4A receptors bind as monomers to the DNA element NBRE or as homo- or heterodimers to the DNA element NuRE. The experiments shown were performed with NuRE. First, we verified that SUMO NR4A1 mutant maintains basal transcriptional activity, and then we compared its transcriptional activity with that of NR4A1 WT in response to SP. As can be seen in Figure 3A, NR4A1_TriMut showed enhanced transcriptional activity in response to SP signaling. Therefore, SUMOylation appears to be a relevant negative regulator of NR4A1 transcriptional activity.

**Figure 3.**
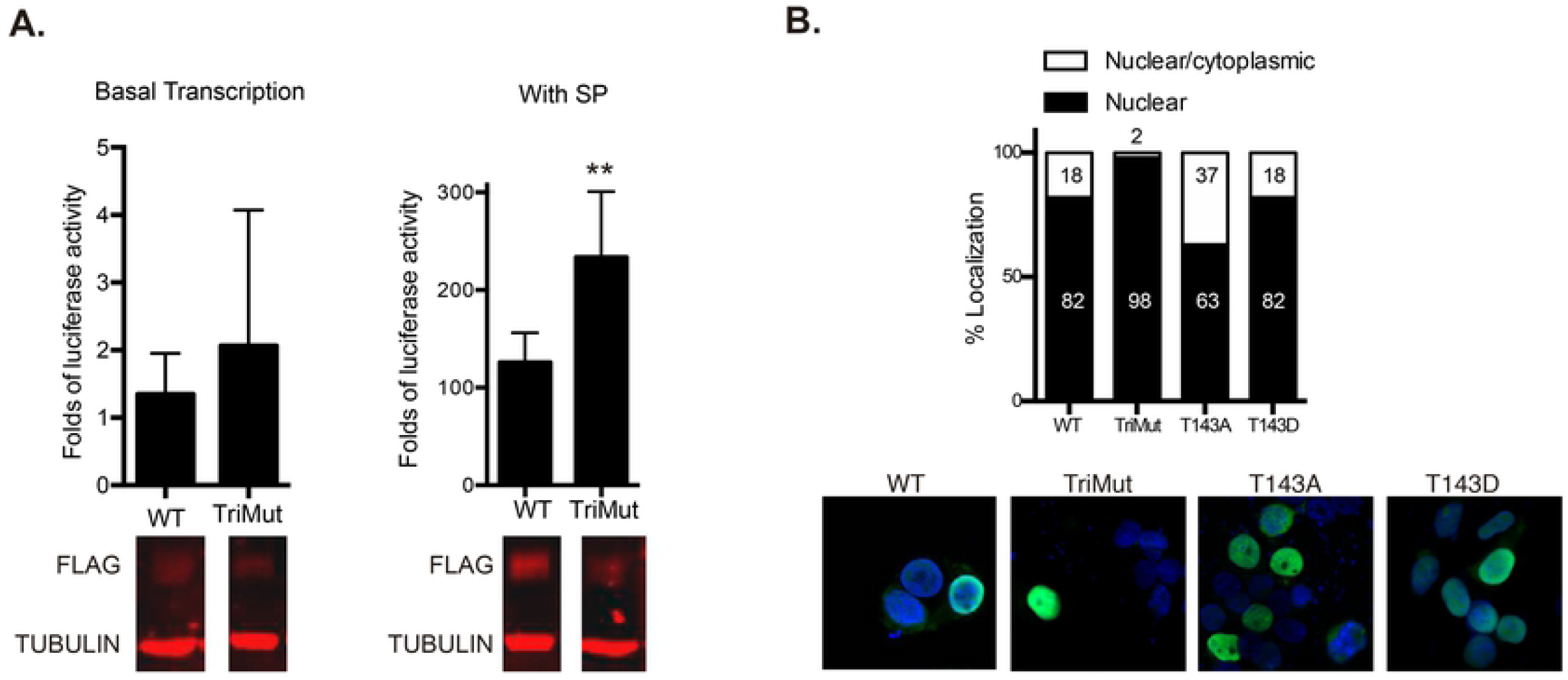
SUMOylation regulates NR4A1 transcriptional activity and its intracellular distribution. **A**, *NR4A1 mutant in Lys residues located in SUMOylation motifs (TriMut) have increased transcriptional activity compared to wild type in response to SP*. Cells were transfected with a reporter containing Luciferase cDNA under the control of NuRE/POMC and the indicated plasmids to assess basal transcriptional activity (left panel). Then cells were co-transfected with NK_1_R and treated or not with SP for 3 hr. TriMut showed enhanced transcriptional activity in response to SP and at a higher level than NR4A1 WT. A mean of 5 independent experiments (each with duplicate wells) is plotted. Error bars represent the standard error. ** p<0.01; n=5. A representative Western blot showing the level of expression of each construct, with or without SP, is shown below. Both anti-FLAG and anti-Tubulin were incubated simultaneously. **B**, *SUMOylation and phosphorylation regulate NR4A1 intracellular distribution*. Cells were transfected with indicated plasmids and the intracellular localization of the proteins was determined by immunofluorescence to detect NR4A1 (green). Two-hundred cells from two independent experiments for each construct were counted. The percentage of cells with only nuclear, or nuclear-cytoplasmic localization of NR4A1, is plotted. Representative confocal microscopy images are shown below; nuclei were stained with DAPI (blue).

NR4A1 can have both nuclear and cytoplasmic functions(15); during SP/NK_1_R–induced autosis around 15% of the cells showed NR4A1 also in the cytoplasm (13). To assess whether SUMOylation affects NR4A1 localization, we analyzed the intracellular distribution of the TriMut. In accordance with the higher level of transcriptional activity we observed, the TriMut was more frequently retained in the nucleus. We also tested the mutant in the target of phosphorylation T143, as it has been previously reported that phosphorylation of NR4A1 also modulates its localization, although in response to other kinases and phosphorylated in other sites (38). We compared NR4A1_T143A and NR4A1_T143D, phosphorylation-defective and threonine phosphorylation-mimic (D mimics the negative charge provided by phosphorylation) mutants, respectively. While most of the cells have NR4A1 restricted to the nucleus, again we observed a small proportion (18%) of cells with NR4A1 also in the cytoplasm (Figure 3B). It appears then than SUMOylation regulates the exportation of NR4A1 out of the nucleus, while phosphorylation might be necessary for retaining NR4A1 in the nucleus. A fine regulation of combined PTMs could determine the intracellular distribution of NR4A1.

### NR4A1 SUMOylation is necessary for SP-induced autosis

We hypothesized that the signaling pathway activated upon NK_1R_ activation by SP binding triggers SUMOylation of NR4A1, conferring upon it the ability to induce autosis instead of apoptosis or proliferation, among other processes. To inhibit NR4A1 SUMOylation we expressed GAM1, a viral protein that inhibits SUMOylation by promoting SAE1/SAE2 and UBC9 degradation (39). As shown in Figure 4A, in the presence of GAM1, but not of an inactive GAM1 mutant, the amount of SUMOylated NR4A1 was reduced. Supporting the notion that SUMOylation increases its basal stability, the total amount of NR4A1 was also reduced in the presence of GAM1. More significantly, *Gam1* expression completely prevented SP/NK_1_R–induced autosis (Figure 4B).

**Figure 4.**
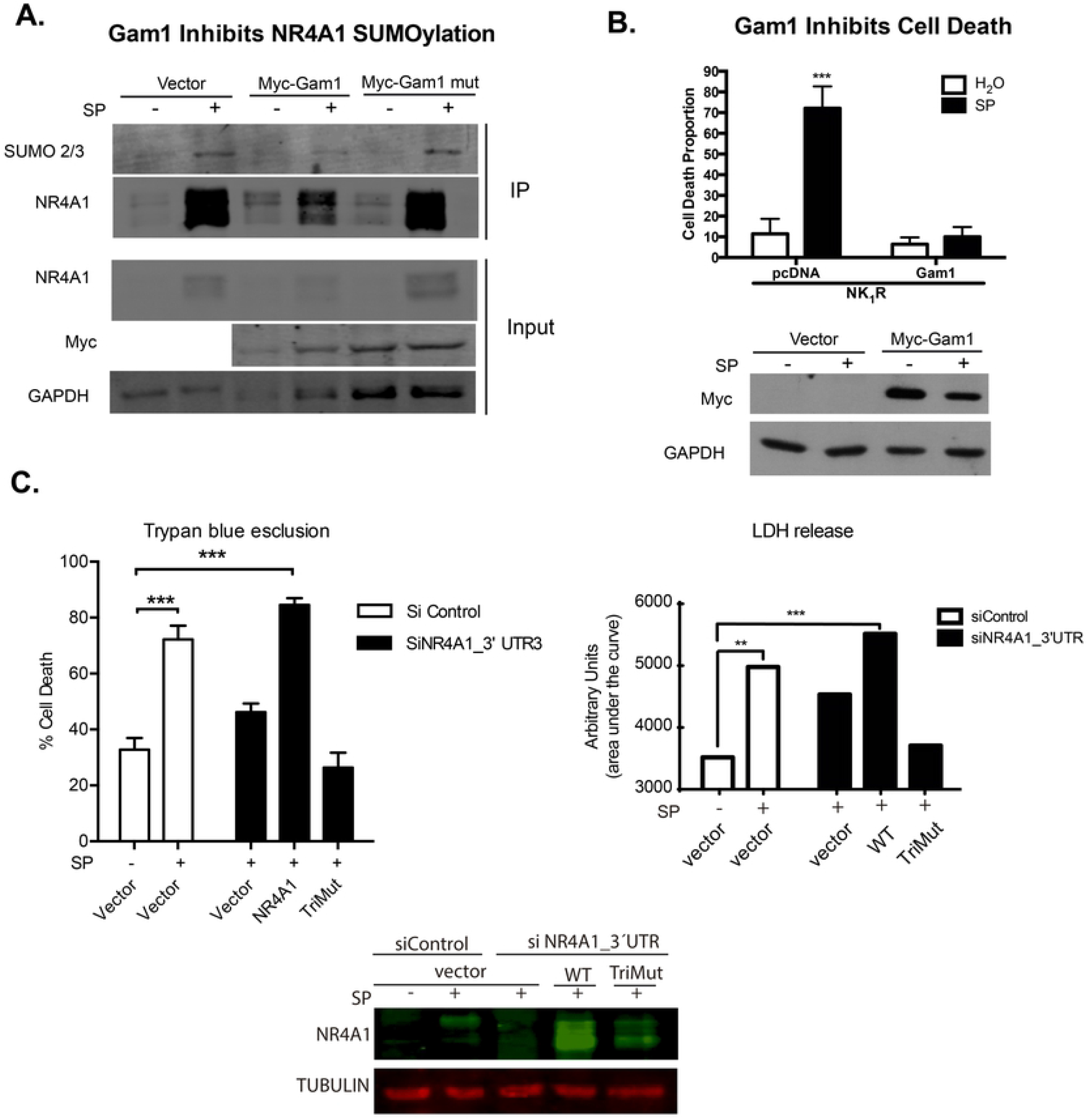
NR4A1 SUMOylation is necessary for SP-induced autosis. **A**, *The viral protein Gam1 reduced SUMOylation of NR4A1*. Cells were transfected with NK_1_R expression vector and the indicated plasmids, and were treated or not with SP for 3 hr. Then, NR4A1 was immunoprecipitated and developed with anti-SUMO2/3 or antiNR4A1. Notice that only in the presence of GAM1, and not with inactive mutant GAM1, there was a reduction in SUMOylated NR4A1. GAPDH was detected in total extract (input) as a reference of initial similar amount of protein. **B**, *Inhibiting SUMOylation by the expression of the viral protein GAM1 prevents SP-induced autosis*. Cells were transfected with NK_1_R expression vector and an empty vector or *Gam1* expression vector and treated or not with SP for 24 hr. Cell death was estimated by Trypan blue exclusion. A mean of three independent experiments is plotted. Error bars represent the standard deviation. *** p<0.0001 2way ANOVA. The Western blot below shows the expression of *Myc-Gam1 or Myc-Gam mutant*. **C**, *SUMOylation defective mutant (TriMut) is not able to induce autosis in response to SP*. Cells were co-transfected with NK1R expression vector and either NR4A1 WT, TriMut or empty vector as indicated. Cells were also transfected with a control siRNA targeting a viral sequence not present in mammals or a siRNA targeting the 3’ UTR, only present in endogenous mRNA. Cells were exposed or not to SP for 24 hr and cell death was estimated by Trypan blue exclusion or LDH activity released. Every experiment was performed in triplicate and averaged. The mean of three independent experiments is plotted. Error bars represent the standard deviation. ***, p<0.0001. Total protein extracts were obtained from replica wells taken at 3hr after SP addition to estimate the content of NR4A1 by WB. Tubulin was detected as a loading reference.

Finally, to confirm whether SUMOylation of NR4A1 is indeed necessary to mediate SP-induced autosis, we tested the ability of TriMut to induce autosis in response to SP. To eliminate endogenous expression of NR4A1 that could mask the mutant phenotype, we silenced it by targeting small interfering RNAs to the 3’ untranslated region of RN4A1 mRNA, which is absent in the NR4A1 expression vector. As expected, silencing the expression of endogenous NR4A1 reduced cell death in response to SP, which was restored by the expression of NR4A1 WT but not by the expression of TriMut (Figure 4C). Therefore, SUMOylation of NR4A1 is necessary for SP/NK_1_R–induced autosis.

## Discussion

NR4A receptors regulate multiple processes such as metabolism, proliferation, migration, apoptosis, DNA repair and autophagy. Accordingly, NR4A receptors are involved in several pathological processes like cardiovascular diseases, diabetes, atherosclerosis and neurodegeneration (40). Interestingly, NR4A1 expression is also induced by caloric restriction, and a reduction in expression of at least NR4A2 accompanies human aging (2). Since DNA damage accumulates with aging, perhaps the reduced expression of NR4A is a contributory factor. Intriguingly, in cancers from different origins, such as melanoma, breast, colon and pancreas, both pro- and anti-tumorigenic activities have been described for NR4A family proteins (7, 41). Indeed, pharmacological regulation of NR4A activity has been proposed to not only counteract aging, including cognitive decline (40), but also cancer and metabolic diseases. Therefore, understanding the mechanisms that regulate NR4A function is an active area of research.

SUMO modification affects numerous cellular processes overlapping with those described for NR4A1 function. Disruption of SUMOylation affects differentiation of cells representative of the three germ layers: endoderm, ectoderm and mesoderm (42). SUMOylation also affects DNA repair and stability, as it acts as a master organizer of protein complexes with functions in chromatin remodeling, double-strand break repair and ribosome biogenesis (36).

In the present work, we demonstrated that NR4A1 is SUMOylated during SP–induced autosis. Specific mutants of NR4A1 with reduced SUMOylation have decreased basal stability but yet responded to SP signaling increasing its stability, as well as its transcriptional activity. This latter effect could be due to additional SUMOylation sites of NR4A1 since there is still a weak SUMO signal in the triple mutant. A recent more comprehensive analysis including many SUMO substrates have shown that the E residue is preferred over D in the consensus motif (43), so there could be additional SUMOylation motifs in the NR4A1 sequence. Even though RNF4 can attach ubiquitin to SUMO-modified NR4A1 in response to signals such as PMA and target it for proteasomal degradation (26), the ubiquitination sites for basal NR4A1 degradation seem to be different to Lys 102, 558 and 577, since when those residues were substituted by arginine the stability of TriMut was reduced. Clearly NR4A1 can have different ubiquitination sites in different cellular contexts. For example, during inflammation, ubiquitinated NR4A1 is not send to proteasomal degradation but functions as a label of damaged mitochondria, interacting with the autophgic receptor p62/SQSTM1 for their engulfment and elimination by mitophagy (44). Further experiments will be needed to understand the combinations of PTMs that regulate NR4A1 half-life, intracellular localization and interactors, which render specific functions in different contexts.

Finally, we show that the ability of NR4A1 to induce autosis is impaired when SUMOylation is reduced. Autosis could be triggered by specific molecular interactions of SUMOylated NR4A1 in the cytoplasm or in the nucleus. Interestingly, NR4A1_TriMut (NR4A1_K102,558,577R) showed a nuclear localization in 98% of the cells. It is possible that SUMOylation in K558 is required for cytoplasmic NR4A1 localization, as K558 is part of the third nuclear export signal (LLGKLPELRTL) located in the LBD (45). Therefore, NR4A1_TriMut’s inability to induce autosis might relate to a reduction in a cytoplasmic function or an alteration in a NR4A1 transcriptionally-regulated pathway that induces autophagy. The present work encourage further experiments to underscore the molecular mechanisms by which NR4A1 and SUMOylation influence the induction of autosis, or by the repression of a target gene. This knowledge is particularly relevant, since NR4A1 expression is also induced in response to ischemia (24) and kainic acid-triggered seizures (25), situations where Substance P mediates neuronal cell death. Screening for small molecules able to inhibit NR4A1 SUMOylation, or inducing its deSUMOylation, would potentially contribute to treatments that prevent or reduce neuronal cell death. From a different point of view, since both NR4A(4, 5) and SUMO (46) regulated pathways have functions in memory, SUMOylation of NR4A1 could have a role in cognitive activities. Altogether, understanding the molecular regulation of NR4A1 function has a potential impact in biomedicine.

## Acknowledgements

We acknowledge the technical assistance of M.C. Concepcion Valencia and Dr. Beatriz Aguilar, as well as the computational support from Ana María Escalante and Francisco Pérez and maintenance of equipment from Aurey Galván and Manuel Ortínez.

## Author contribution

GZG, GMH, MRSC, WVG, APPR and CA contributed to the investigation and formal analysis; MEA and LC contributed with resources and supervision; CW contributed to visualization; SCO contributed with conceptualization, funding acquisition, project administration, resources, supervision and writing original draft. All authors reviewed and edited the manuscript. Data in this work are part of GZG and GMH dissertation thesis in the “Posgrado en Ciencias Bioquímicas de la Universidad Nacional Autónoma de México”. WVG and APPR were undergrad students of Facultad de Ciencias, UNAM.

## Abbreviations list

NR4A: Nuclear receptor group family A
NR4A1: Nuclear receptor group 4 family A member 1 (Nur77, NGF1B, TR3, etc.)
NR4A2: Nuclear receptor group 4 family A member 2 (Nurr1, NOT1, etc)
NR4A3: Nuclear receptor group 4 family A member 3 (Nor1, MINOR)
SUMO: small ubiquitin-like modifier
SP: Substance P
NK_1_R: Neurokinin 1 Receptor
TAD: Trans-Activation Domain
DBD: DNA Binding Domain
LBD: Ligand Binding Domain
PTM: Post Translational Modifications

